# Fluorescent Antibody Multiplexing with Oligo-Based Combinatorial Labeling

**DOI:** 10.1101/2020.11.06.371906

**Authors:** Madeline McCarthy, Caitlin Anglin, Heather Peer, Sevanna Boleman, Stephanie Klaubert, Marc R. Birtwistle

**Affiliations:** Department of Chemical and Biomolecular Engineering, Clemson University, Clemson SC 29634, USA

## Abstract

Fluorescent antibodies are a workhorse of biomedical science, but fluorescence multiplexing has been notoriously difficult due to spectral overlap between fluorophores. We recently established proof-of-principal for fluorescence Multiplexing using Spectral Imaging and Combinatorics (MuSIC), which uses combinations of existing fluorophores to create unique spectral signatures for increased multiplexing. However, a method for labeling antibodies with MuSIC probes has not yet been developed. Here, we present a method for labeling antibodies with MuSIC probes. We conjugate a DBCO-Peg5-NHS ester linker to antibodies, a single stranded DNA “docking strand” to the linker, and finally, hybridize two MuSIC-compatible, fluorescently-labeled oligos to the docking strand. We validate the labeling protocol with spin-column purification and absorbance measurements, which show a degree of labeling of ~9.66 linker molecules / antibody. We demonstrate the approach using (i) Cy3, (ii) Tex615, and (iii) a Cy3-Tex615 combination as three different MuSIC probes attached to three separate batches of antibodies. We incubated MuSIC probe-labeled antibodies with protein A beads to create single and double positive beads that are analogous to single cells. Spectral flow cytometry experiments demonstrate that each MuSIC probe can be uniquely distinguished, and the fraction of beads in a mixture with different staining patterns is accurately measured. The approach is general and might be more broadly applied to cell type profiling or tissue heterogeneity studies in clinical, biomedical, and drug discovery research.

## Introduction

Ultraviolet-to-infrared fluorescence is a bedrock of experimental science, particularly the biomedical sciences. However, multiplexing—the simultaneous analysis of multiple fluorophores in a single sample, is severely limited by spectral overlap^1–4^, where excitation and/or emission spectra of fluorescent probes share broad wavelength domains. Spectral overlap limits most standard fluorescence assays to 2-4 readouts at a time. Yet, many applications would benefit from increased fluorescence multiplexing capabilities; one example is cancer. Tumor heterogeneity is multi-dimensional including spatial variation in cell type, driver mutation profiles, protein expression, and oxygen/metabolic gradients^5–10^. As a result, there are hundreds of markers that have an impact on a tumor’s evolution, fitness, and drug sensitivity ^5,11,12^.

Current sequencing methods can reach high levels of multiplexing and have been used in cancer diagnosis and prognosis^13–15^. Yet, the now somewhat standard biopsy- or homogenized tissue-based deep DNA or mRNA sequencing, and now increasingly single-cell sequencing^16–18^, largely do not allow for spatial resolution. However, some recent sequencing-based methods can provide spatial *in situ* data^19–22^. Sequential fluorescence in situ hybridization (seqFISH+) is capable of transcriptome-wide imaging in single cells, but has challenges in scaling to large numbers of cells or large areas of tissue sections. Slide-seq, alternatively, made mRNA sequencing compatible with tissue section imaging over large spatial scales with ~10 um resolution^23^. Although powerful advances, such sequencing methods cannot yet fully capture the heterogeneity of tissue samples, which includes single and subcellular resolution and molecules other than mRNA (i.e. DNA, proteins, post-translational modifications, etc.). Antibody-based imaging, on the other hand, can access multiple molecule types at single and subcellular resolution while also spanning physiologically relevant length scales. Therefore, increased antibody multiplexing capabilities remains highly complementary to these sequencing-based methods.

There have been many recent advances for increased antibody-based multiplexing with single cell and subcellular spatial resolution, most of which use standard “filter-based” instrumentation that robustly allow imaging 2-4 fluorescence colors simultaneously. A widely adopted strategy is repeated rounds of staining, imaging, and bleaching of fluorophores^24–27^. By performing multiple cycles of 2-4 color imaging, these methods drastically increase fluorescent multiplexing capabilities (up to 60 analytes). Multiplexed fluorescence microscopy (MxIF) was the first but requires proprietary and expensive equipment / reagents^24^. Cyclic Immunofluorescence (CyCIF) is similar in principal but uses inexpensive reagents and standard equipment^26,28^. Similar to MxIF and CyCIF, Iterative indirect immunofluorescence imaging (4i) uses cycles of imaging, but leverages fluorophore-conjugated secondary antibodies rather than fluorophore-conjugated primary antibodies as in the above techniques, allowing use of “off-the-shelf” primary antibodies^25^. Another method that makes use of staining and bleaching cycles is co-detection by indexing (CODEX)^27^, but it differs from the above methods as it uses DNA-conjugated antibodies and sequencing-like methods to multiplex. While these cyclic methods have significantly expanded multiplexing capability, a main limitation is the number of rounds of imaging that are possible before sample degradation begins to occur. Additionally, the length of time each round takes to complete, multiplied by the number of rounds, can make these methods excessively time consuming.

Another way to achieve higher degrees of antibody multiplexing is by labeling antibodies with isotopically pure rare earth metals, such as in imaging mass cytometry (IMC) ^29^ and multiplexed ion beam imaging (MIBI) ^30^. IMC and MIBI can respectively image 32 and 40 analytes simultaneously from a tissue sample. The use of mass spectrometry for quantification makes these techniques easier to multiplex compared to ones that use fluorescence, as they are not limited by spectral overlap. However, these methods use a laser or ion beam to ablate the sample, destroying the sample and preventing further analysis or use, including cyclic methods as above. Additionally, the specialized equipment and reagents required for these techniques can be more expensive than standard fluorescence microscopes and antibodies, making them not as widely available.

The fluorescence-based techniques that were previously described use “filter-based” imaging that lumps emission wavelengths together and thus restricts multiplexing to 2-4 channels, but some have instead used spectral imaging that measures emission intensity with much finer wavelength resolution. Fluorescence emission follows the principle of linear superposition, meaning that the emission spectra of a mixture of fluorophores can be cast as a sum of contributions from individual probes using a matrix equation. Solving this matrix equation for the levels of individual probes, given the spectra of the mixture and each isolated probe, is called unmixing. These “hyperspectral” techniques have been used to image up to seven analytes simultaneously in tissue sections^3,31–34^. CLASI-FISH (combinatorial labeling and spectral imaging - fluorescence in situ hybridization), which builds upon traditional spectral imaging, classifies up to 15 microbe types using probe combinations^35^. One constraint of CLASI-FISH is that probes must be spatially segregated for demultiplexing. Spectrally resolved fluorescence lifetime imaging microscopy (sFLIM)^36^ combines spectral imaging with fluorescence lifetime information and can multiplex nine antibodies simultaneously.

We recently developed an approach called Multiplexing using Spectral Imaging and Combinatorics (MuSIC), which leverages currently available fluorophores along with the power of combinatorics to increase the number of available probes for simultaneous staining^37^. Förster resonance energy transfer (FRET), a phenomenon whereby a higher energy fluorophore donates energy to a lower energy fluorophore^38^, is central to creating MuSIC probes with unique spectral signatures. MuSIC probes are created using FRET-producing fluorophore combinations, which results in a probe emission spectrum that is linearly independent from that of the individual fluorophores that make up the combination, enabling unmixing. Our previous work, based on simulation, suggested that MuSIC may increase simultaneous fluorescence multiplexing capabilities ~4-5 fold^37^. Proof-of-principal experimental studies that focused on a small range of excitation wavelength space have shown that nine MuSIC probes can be accurately unmixed, which should increase when the full range is used. Moreover, MuSIC is compatible with cyclic imaging methods, which would allow more analytes to be measured per cycle, increasing multiplexing capabilities even further. MuSIC differs from CLASI-FISH in that it is not limited by spatial segregation.

Although MuSIC has potential to increase fluorescence multiplexing, methods to conjugate MuSIC probes to antibodies have not yet been developed. Our previous work showed that standard primary-amine-based conjugation of two fluorophores to the same antibody does not produce a high enough FRET efficiency to create robust MuSIC probes^39^. Here, we report a fluorescent oligo-based labeling approach to conjugate MuSIC probes to antibodies. A DBCO-Peg5-NHS ester molecule (the linker) is used to attach an azide modified oligo (the docking strand) to the antibody. Fluorescent oligos hybridized to the docking strand bring the fluorophores into FRET-compatible distances. Mixtures of antibody-conjugated MuSIC probes using (i) Cy3, (ii) Tex615, and (iii) a Cy3-Tex615 combination were analyzed and accurately unmixed using spectral flow cytometry as a proof-of-principle. These oligo-based MuSIC probes are compatible with the wide range of clinical, biomedical and drug discovery applications that currently use fluorescent antibodies and spectral imaging.

## Methods

### Adding the linker to the antibody

This and the below procedures were developed around labeling 50 mg of IgG but is compatible with scaling up or down. In our case, normal Rabbit IgG (ThermoFisher Cat: 31235) is combined with DBCO-Peg5-NHS Ester (linker; 10 mM in DMSO; Click Chemistry Tools Cat: 1378531-80-6) in 60 molar excess (50 mg of Rabbit IgG and 4.6 mg of linker). This is brought to a volume of 100ml with PBS and allowed to incubate for 30 minutes at 25°C. After incubation, the solution is added to an Amicon Ultra 0.5ml 100 kDa centrifugal filter (Fisher Scientific Cat: UFC5100BK) and spun for 5 minutes at 14,000 x g. The filter is then placed into a new tube and PBS is added to the top of the filter in order to bring the total volume back to 100 ml and is spun again for 5 minutes at 14,000 x g. This wash step is repeated twice (three total). Finally, the filter is flipped upside down and placed in a clean tube and spun for 1 minute at 1000 x g to collect the retentate. The retentate absorbance is measured at 309nm, where the linker strongly absorbs, and 280nm, where the antibody strongly absorbs, using a NanoDrop spectrophotometer (Thermo Scientific).

### Adding the docking strand to the antibody-linker conjugate

The docking strand (Integrated DNA technologies-Table 1) is added to the antibody-linker retentate from the previous step in 6 molar excess to the original amount of antibody (2 nmoles of docking strand). The volume is brought up to 100 ml with PBS and incubated at 4°C overnight. The sample is then placed in an Amicon Ultra 0.5ml 100 kDa centrifugal filter and spun for 5 minutes at 14,000 x g. Once this spin is completed, the filter is placed into a new tube and PBS is added to the top of the filter in order to bring the total volume back to 100 ml and is spun again for 5 minutes at 14,000 x g. This wash step is repeated twice. Finally, the filter is flipped upside down and placed in a clean tube and spun for 1 minute at 1000 x g to collect the retentate. The retentate absorbance is measured at 309nm and 280nm as above, and also at 260nm, where the docking strand strongly absorbs, using a NanoDrop spectrophotometer (Thermo Scientific).

**Table 1:**
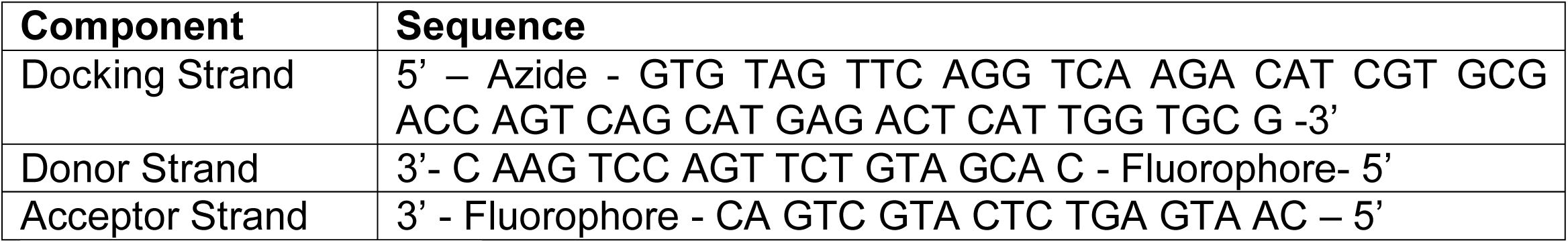
Oligo Sequences

### Degree of Labeling

To generate calibration curves for concentrations of the antibody, linker, and docking strand, absorbance measurements are taken using a NanoDrop spectrophotometer (Thermo Scientific) for known concentrations of the antibody, linker, and docking strand at 309, 260, and 280nm. Five-point, 2-fold serial dilutions were used to generate samples for the calibration curve. A least squares line of best fit (MATLAB) is generated to estimate absorbance extinction coefficients based on Beer’s law (1) for each component at 309, 280, and 260nm.

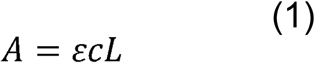

Here, *A* is the absorbance of the solution, *e* is the extinction coefficient, *L* is the length of path traveled by light (1 mm), and *c* is the concentration of the solution. From here, a system of three simultaneous equations is solved in Matlab using the function `vpasolve` to estimate concentrations of the antibody (a), linker (l), and docking strand (DS) given absorbance measurements at 309, 280 and 260 nm from a mixture (M).

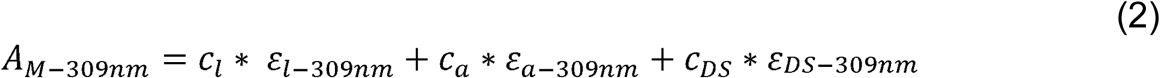

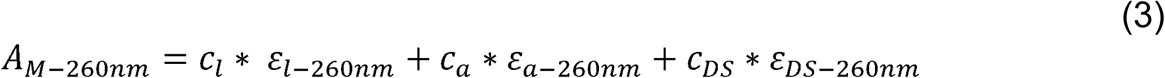

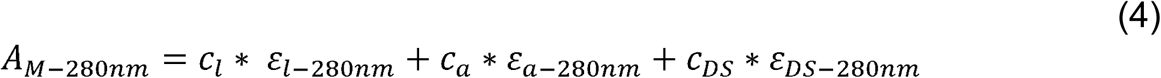

Degree of labeling for linker to antibody could be calculated from the above estimated concentrations. However, due to the nature of the spin column-based separation, some unreacted linker will remain. This amount of residual linker can be calculated based on mole balance and we used this calculation to correct the degree of labeling as follows, where *n* is the number of washes

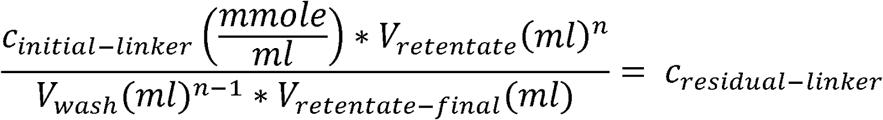

The concentration of the residual linker is subtracted from the calculated linker concentration to determine the concentration of linker that is attached to the antibody in the retentate.

### Adding the donor and acceptor strands to the antibody-linker-docking strand conjugate

A 25 bp oligo with a 5’ fluorophore modification (donor strand) and a 25 bp oligo with a 3’ fluorophore modification (acceptor strand) (each 100 mM in water, Integrated DNA technologies) are added in equimolar amounts (2 nmoles each) to the antibody-linker-docking strand retentate and brought up to 100 ml with PBS. Sequences are shown in Table 1. This solution is allowed to incubate for 15 minutes at 25°C in the dark. When testing the necessity of the linker, the donor strand with an Atto 488 modification was added to the antibody-linker-docking strand retentate. To make the different probes, Probe 1 consists of equimolar amounts of the donor strand with a Cy3 modification and the acceptor strand with a Cy3 modification (each 2 nmoles), Probe 2 consists of equimolar amounts of the donor strand with a Tex615 modification and the acceptor strand with a Tex615 modification (each 2 nmoles), and Probe 3 consists of equimolar amounts of the donor strand with a Cy3 modification and the acceptor strand with a Tex615 modification (each 2 nmoles).

### Choosing donor and acceptor pairs

To test the donor and acceptor fluorophore pair of Cy3 and Tex615, 4 samples are created. (1) A 25 bp oligo with a 5’ Cy3 modification (donor strand) and a 25 bp oligo with a 3’ Tex615 modification (acceptor strand) (each 100 mM in water) are added in equimolar amounts (0.2 nmoles), (2) The donor, acceptor, and docking strands are added in equimolar amounts (0.2 nmoles), (3) 0.2 nmoles of the donor strand, (4) 0.2 nmoles of the acceptor strand. All samples are brought to 50 ml with PBS. The samples (oligos in solution) are loaded into a black 96 well plate (Fisher Scientific Cat: 655900) and fluorescence emission spectra assayed with a Synergy MX microplate reader (Biotek). Parameters are set to a slit width of 9nm, a 10 second shake prior to reading, taking readings from the top, and excitation wavelength of 488 nm. The emission start ranges are 50nm greater than the excitation wavelength.

### Attaching labeled antibodies to protein A dynabeads

The MuSIC-probe labeled antibodies from above are suspended in 200 ml of 0.02% (2 ml/10ml) Tween 20 (Fisher Scientific Cat: 9005-64-5) in PBS and added to 50 ml of protein A dynabeads (Fisher Scientific Cat: 10 001 D—33 mg of initially added IgG; 100 mg batch makes three incubations). For making double positive beads, both probes are simultaneously added to 50 ml of protein A dynabeads. This solution is allowed to incubate for 10 minutes with rotation in the dark. After incubation, the solution is placed on a magnet, the supernatant is removed, and the bead-antibody complex is resuspended in PBS with Tween 20. The solution is then placed back on the magnet and the supernatant is again removed and is resuspended in PBS.

### Analyzing probe mixtures using Cytek Aurora flow cytometer

Mixtures of bead-conjugated probes are analyzed using a Cytek Aurora spectral flow cytometer with 488nm and 635nm excitation lasers. First, beads with single probes are assayed as reference controls. The events to record is set to 5,000, the stopping time is set to 10,000 sec, and the stopping volume is set to 3,000 ml. For samples containing mixtures of bead types or double-positive beads, the events to record are set to 15,000, the stopping time is set to 10,000 sec, and the stopping volume is set to 3,000 ml. Once mixtures have been analyzed, the SpectroFlo software (Cytek) is used first gate single beads by gating over the population of beads with the lower forward and side scatter area, and then to unmix and report (i) the amount of each probe on every bead that was analyzed and (ii) the fraction of each bead type in each mixture of bead types.

## Results

### Probe design and labeling process

A fundamental component of the Multiplexing using Spectral Imaging and Combinatorics (MuSIC) approach is that combinations of fluorophores exhibiting FRET create a unique emission spectrum that is linearly independent from that individual fluorophores in the combination. Thus, creating MuSIC probes on antibodies requires combinations of fluorophores to be stably associated with antibodies with spatial proximity sufficient for FRET. To achieve this, we started from a prior description of antibody-oligo labeling^40^ (Figure 1a). First, a DBCO (dibenzocyclooctyne)-PEG5-NHS ester molecule (referred to as the linker) is attached to the antibody. The NHS ester group at the end of the linker reacts with available NH_2_ groups on the surface of the antibody. From here, a 55 bp DNA oligo with a 5’ azide modification (referred to as the docking strand) is added to the complex. The azide reacts with the DBCO group of the linker via copper-free click chemistry, creating an antibody-linker-docking strand conjugate. The PEG5 group is included in the linker to increase water solubility of the DBCO group and provide space between the antibody and the docking strand^40^. Finally, 25 bp oligos with 5’ or 3’ fluorophore modifications (referred to as the donor and acceptor strands, respectively) are added to the antibody-linker-docking strand conjugate solution. When the donor and acceptor strands hybridize to the docking strand, the two fluorophores are in close physical proximity to enable FRET. The final product of these reactions should be an antibody labeled with a MuSIC probe (Figure 1b).

**Figure 1:**
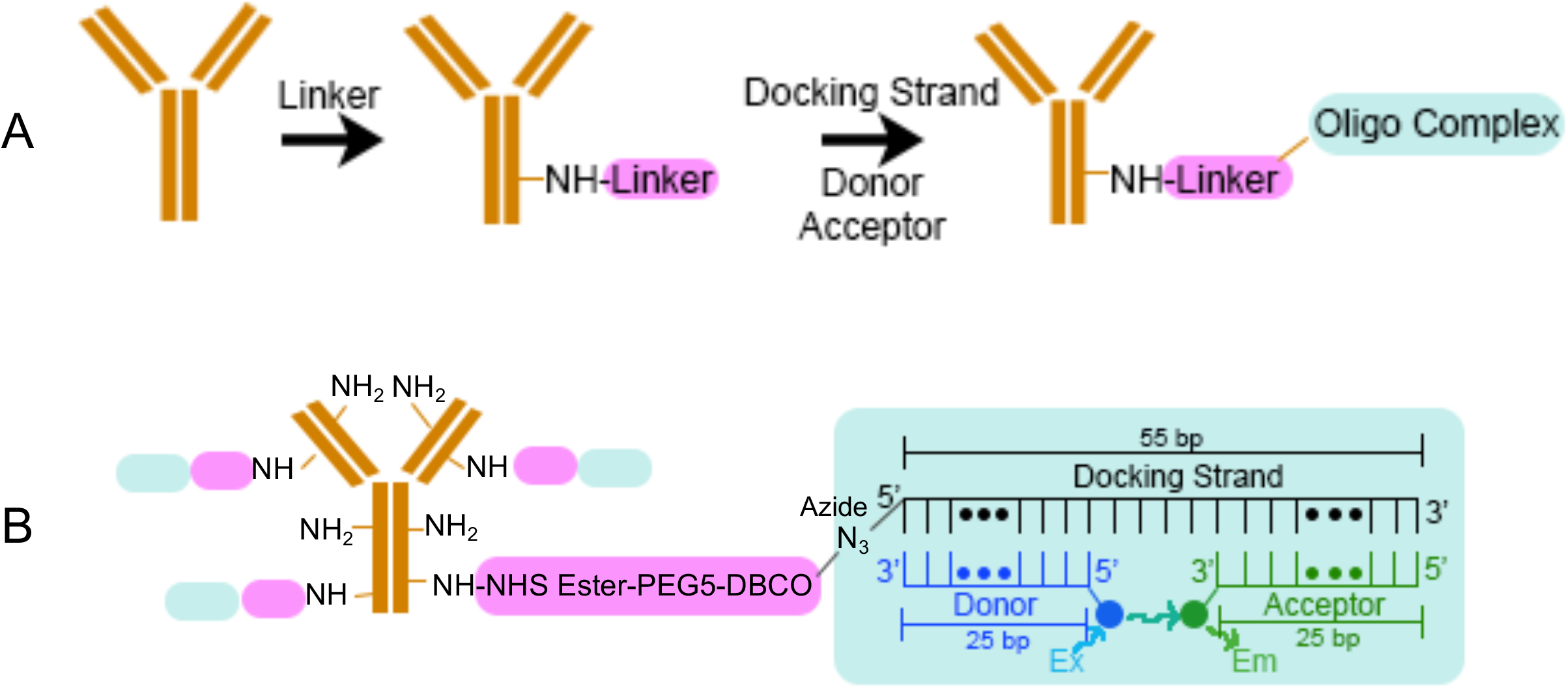
Labeling antibodies with oligo-based MuSIC probes. (A) Labeling schematic for MuSIC probes. First the linker is added to the antibody by reacting the NHS ester on the linker with the NH_2_ group on the antibody. Then the docking strand is added and reacts with the linker via copper-free click chemistry. Lastly, the donor and acceptor strands are annealed to the docking strand to form the oligo complex. (B) Final product of labeling process. The linker can attach to the antibody at multiple NH_2_ sites, allowing an increased degree of labeling.

### Attaching the linker to the antibody

We developed the protocol around labeling 50 mg of IgG, although it is scalable in either direction. The linker is added to the antibody in 60 molar excess, as the linker will react with multiple free amine sites on the surface of the antibody, and the extent of reaction is not certain, but it is desired to maximize degree of labeling. After incubation, unattached linker needs to be separated from the antibody-linker conjugate. To do this, we used Amicon Ultra 100 kDa molecular weight cut off (MWCO) filters (Figure 2a). The antibody has a molecular weight of ~150 kDa and the linker has a molecular weight of 0.7 kDa, so once the solution is spun and washed, any linker that does not attach to the antibody will freely flow through the column (Figure 2b). In order to verify that unattached linker was removed, retentate absorbances were measured at 309 nm, where the linker strongly absorbs^40^ (Fig. S1), for samples containing the antibody alone, the linker alone, and then antibody and linker together. Results show that the linker is predominantly in the retentate only when the antibody is present (Figure 2c). The degree of labeling was estimated to be ~9.66 with a standard error of 1.04 molecules of linker/antibody based on absorbance measurements (see Methods and Figure S1). These results demonstrate that the antibody and linker can stably associate and that unattached linker can be effectively removed from solution.

**Figure 2:**
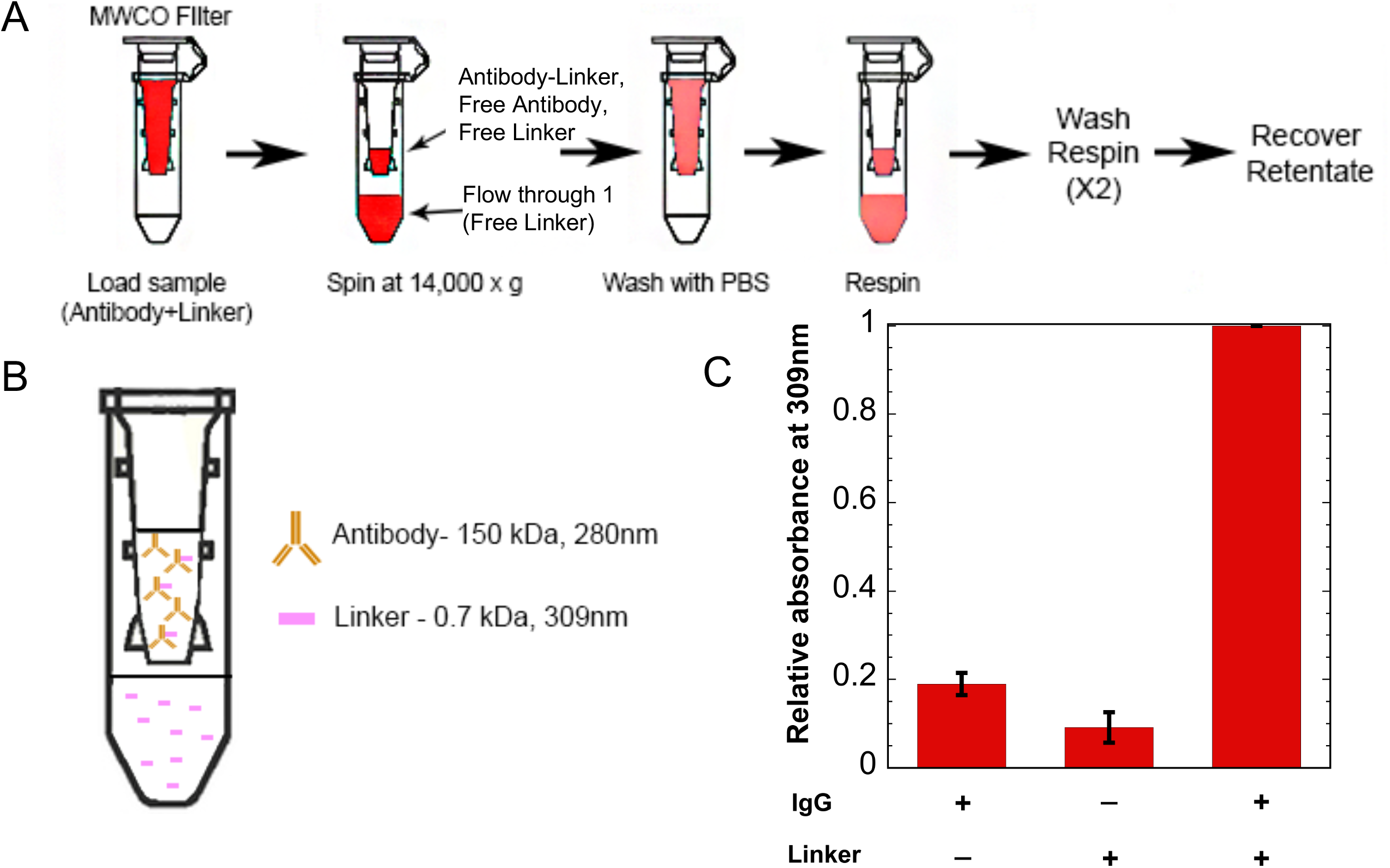
Adding linker to antibody. (A) Separating free linker using 100 kDa molecular weight cut off (MWCO) filters. The antibody and linker are incubated, then the sample is added to the molecular weight cut off filters. The filters are spun to separate unattached linker and then go through a series of washes. Finally, the retentate is recovered. (B) Expected separation of components after spin and wash steps. (C) Retentate absorbances at 309nm. Results show increased signal at 309nm when the linker is in the presence of the antibody.

### Attaching the docking strand to the antibody

The docking strand is added to the antibody-linker retentate from the previous step in 6 molar excess to the antibody to account for multiple labeling sites. After incubation, unattached docking strand needs to be separated from the antibody-linker-docking strand conjugate. Similar to above, we use Amicon Ultra 100 kDa MWCO filters (Figure 3a). Since the docking strand is only 17 kDa, it should freely flow through the columns if it is not attached to the antibody-linker conjugate (Figure 3b). In order to evaluate whether unattached docking strand is removed, retentate absorbances were measured at 260 nm, as this is where the docking strand strongly absorbs. Results show that the docking strand can be seen in the retentate when in the presence of the antibody and the linker, as expected. However, a strong retentate signal was also seen for the docking strand when in the presence of only the antibody, without the linker (Figure 3c). The cause for the strong docking strand signal in the retentate without the linker present is unknown, but before proceeding, we wanted to understand whether the docking strand was stably bound to the antibody without the linker present, or whether it could be removed with further washing via an orthogonal separation mechanism.

**Figure 3:**
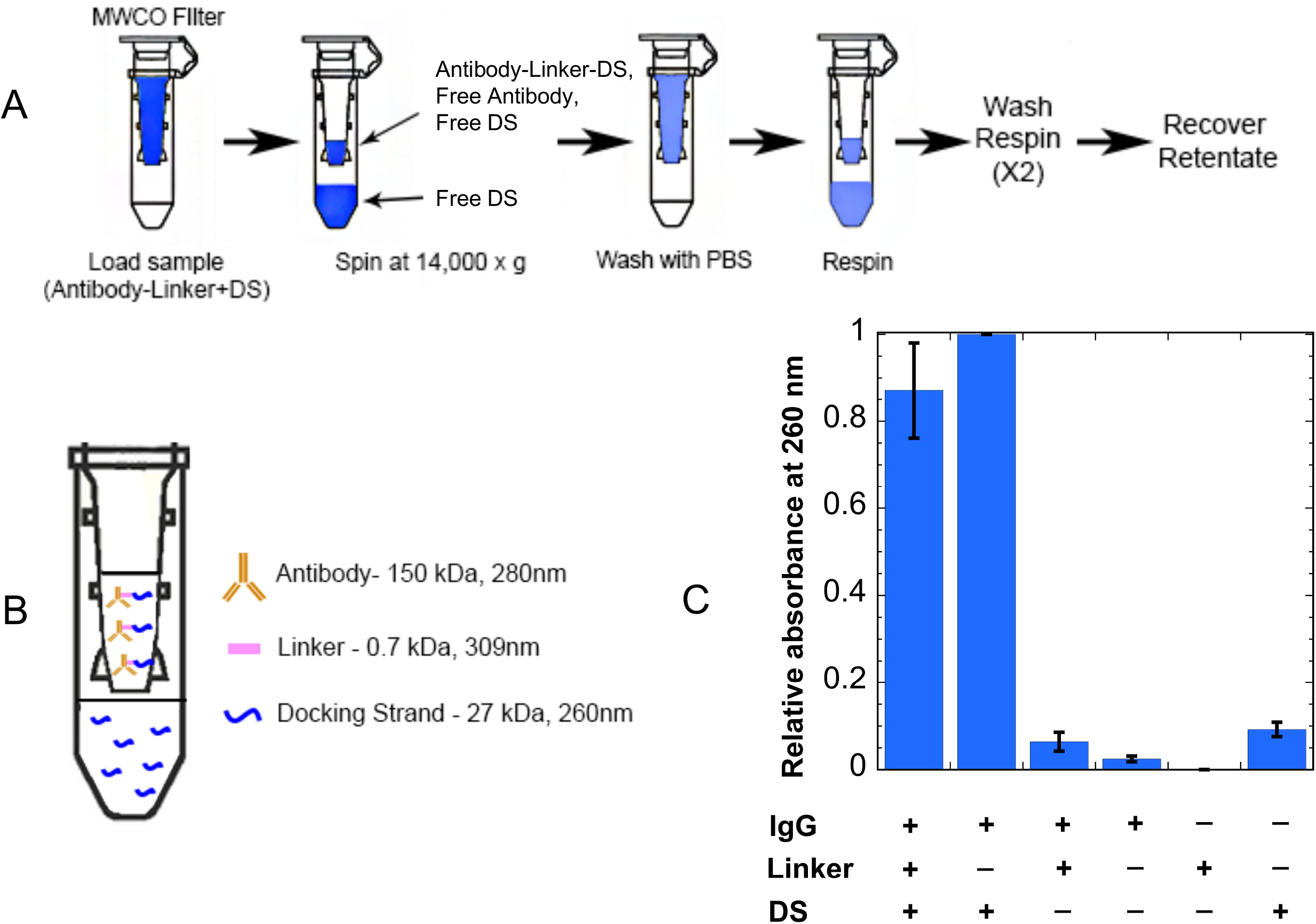
Adding docking strand (DS) to antibody-linker. (A) Separating free DS using 100 kDa MWCO centrifugal filters. The DS and antibody-linker conjugate are incubated, then the sample is added to the molecular weight cut off filters. (B) Expected separation of components after spin and wash steps. (C) Retentate absorbances at 260nm. Results show increased signal at 260nm when the DS is in the presence of the antibody-linker conjugate. An increased signal can also be seen for the case of just the DS and antibody, which is accounted for in later steps.

### The docking strand requires the linker to be stably associated with the antibody

To determine whether the docking strand could stably bind to the antibody without the linker, we used Protein A dynabeads. The beads should strongly and selectively bind to the antibody, and anything attached to the antibody will also be bound to the beads. We created samples with and without linker containing antibody, docking strand, and a donor strand with the fluorophore Atto 488 (for measurement). The supernatant containing any non-stably attached reagents can be removed by washing when the solution is placed on a magnet (Figure 4a). Atto 488 fluorescence was measured to evaluate whether the docking strand could stably associate with the antibody without the linker. The bead-based nature of the experiment precluded reliable absorbance assays as used previously; consequently, we are not able to estimate degree of labeling for the docking strand on the antibody. The fluorescence signal for samples without the linker present was comparable to the signals of the controls where no fluorophore was present, while when the linker was present a significant fluorescence signal was observed (Figure 4b). We conclude that the linker is needed for the antibody to be stably associated with the docking strand, and subsequently, fluorophore-labeled donor or acceptor strands.

**Figure 4:**
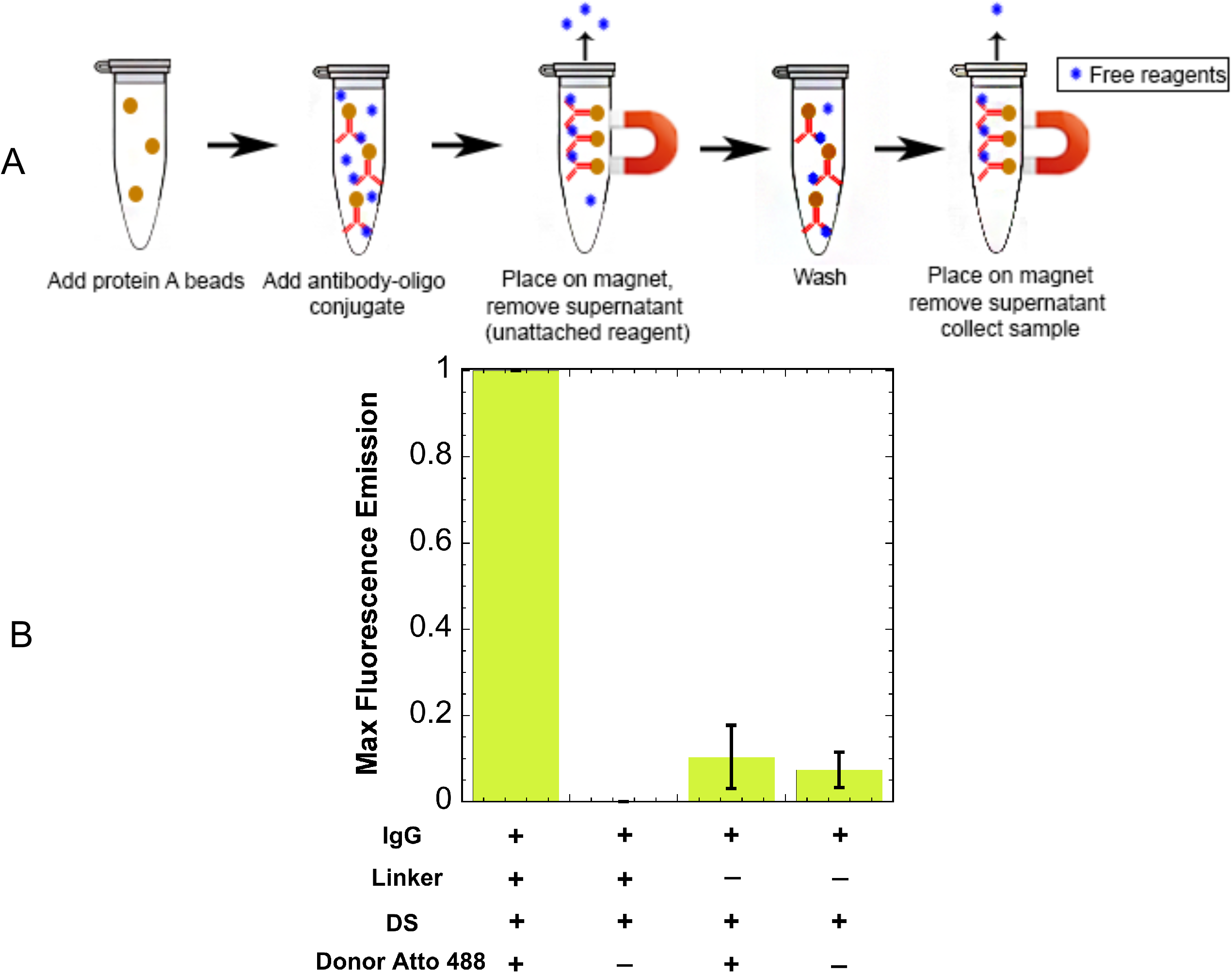
Separating free reagents using protein A beads. (A) The antibody-oligo conjugate is added to the protein A beads and is incubated with rotation for 10 minutes. It is then placed on a magnet, pulling the beads out of solution, and the supernatant containing free reagents is removed. The final product is collected containing the antibody-oligo conjugate. (B) Maximum fluorescence intensity values when excited at 450nm. Results show an increased fluorescence signal for the donor when the linker is added. Without the linker, the fluorescence signal for the donor is the same intensity as the background fluorescence.

### Obtaining a donor and acceptor pair that produce FRET when co-hybridized to the docking strand

As mentioned above, MuSIC probes must have donor and acceptor pairs that exhibit FRET, such that the combination probe has a unique spectral signature. To test if a donor and acceptor pair exhibits FRET, the emission spectra of solutions containing (i) just the donor, (ii) just the acceptor, (iii) the donor and the acceptor, and (iv) the donor and acceptor co-hybridized to the docking strand were analyzed using a plate reader (Figure 5a). We used 488 nm excitation, a common laser line in multiple assay types. The pair of Cy3 (donor) and Tex615 (acceptor) showed a much larger red-shifted emission peak when excited at 488 nm and co-hybridized to the docking strand, indicating strong FRET (Figure 5a). These results showed that this donor and acceptor pair would be a suitable MuSIC probe candidate, i.e. a donor and acceptor strand hybridized to the antibody-linker-docking strand conjugate.

**Figure 5:**
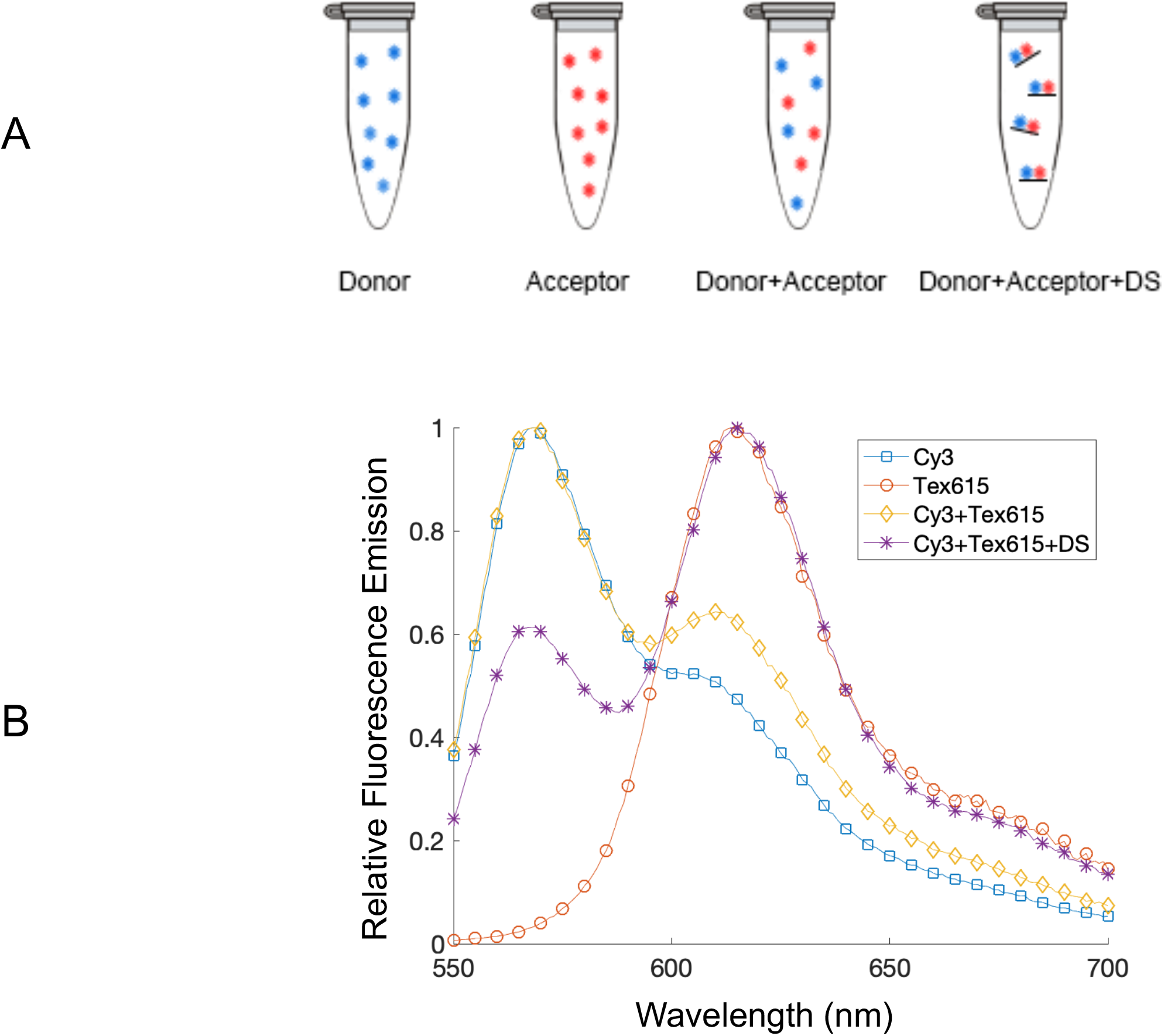
Donor and acceptor fluorophore pair, Cy3 and Tex615. (A) Experimental set up for testing fluorophore combination. We expect only the sample with the DS shows significant FRET. DS: docking strand. (B) Fluorescence emission spectra when excited at 488 nm. An increased acceptor emission peak is seen when the donor and acceptor are annealed to the docking strand, indicating increased FRET.

### Application to flow cytometry for event classification

While there are many potential applications of MuSIC probe-labeled antibodies, we set out to obtain proof-of-principle data using spectral flow cytometry. Namely, we wanted to understand whether we could (i) robustly classify events (i.e. cells) as containing a particular combination of MuSIC probes and (ii) estimate the proportion of events having a particular probe staining pattern (Fig. 6a). This is analogous to cell type classification assays; one widely employed example is to estimate the different abundances of immune cell types in peripheral blood mononuclear cell (PBMC) mixtures^41,42^. Three antibody batches with different probes were created: probe 1-donor Cy3 and acceptor Cy3; probe 2-donor Tex615 and acceptor 615; and probe 3-donor Cy3 and acceptor Tex615. Because Cy3 and Tex615 produce FRET when co-hybridized to the docking strand, probes with this combination of fluorophores can be thought of as a different “color” from the probes with the individual fluorophores of the combination. Once the antibodies with either MuSIC probe 1, 2 or 3 are created, they are incubated with protein A dynabeads to be analyzed using the flow cytometer. Each bead is similar to a single “cell”. One or more antibody type (i.e. with probe 1, 2 or 3) can be conjugated to the same set of beads. For example, incubating beads with two antibody types creates “double positive beads (cells)”. In the following set of experiments, we made single positive (one antibody type conjugated to one bead set) and double positive beads (two antibody types conjugated to one bead set) (Figure 6a). This is related to (i) above. We also make mixtures of these different bead sets. This related to (ii) above. For analysis, we use simple quadrant gates on three bivariate plots to classify beads as negative, single positive, or double positive, and additionally, estimate the proportion of beads that fall into each category (Figure 6b). Populations of single positive beads are observed in R1 and R4, populations of double positive beads are observed in R2, and the negative population of beads is observed in R3 (Figure 6b).

**Figure 6:**
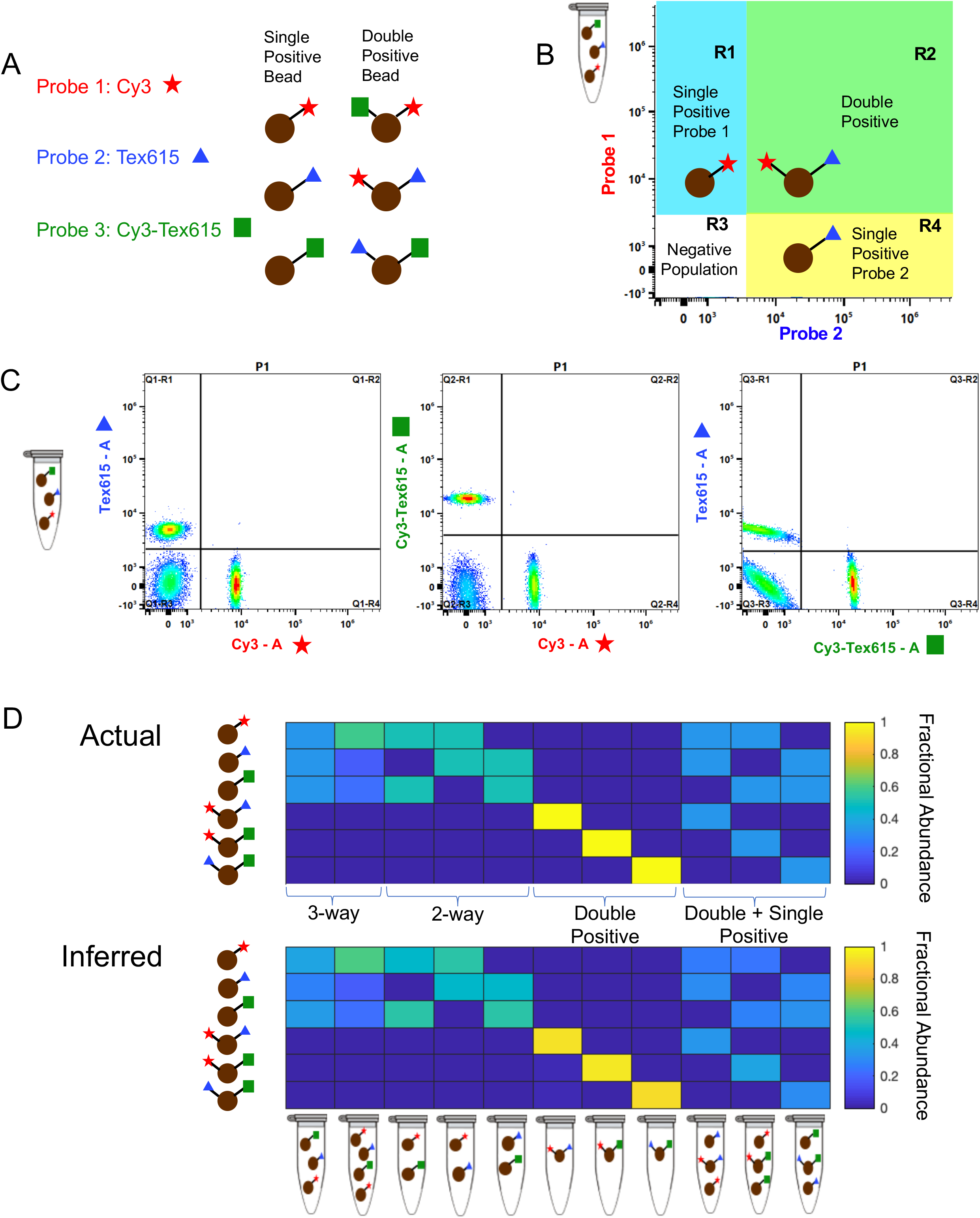
Spectral flow cytometry setup and results. (A) Experimental setup for three-probe mixture ((1) Cy3, (2) Tex615, and (3) Cy3-Tex615) using single-labeled and double-labeled beads. (B) Gating strategy for the populations of beads in the three-probe mixture. (C) Unmixing populations of single-labeled beads in a three-way equimolar mixture of probes Cy3, Tex615, and Cy3-Tex615 using spectral flow cytometry. The plots show unmixing results of Tex615 compared to Cy3 (left), Cy3-Tex615 compared to Cy3 (middle), and Tex615 compared to Cy3-Tex615 (right). (D) Comparing actual amounts of each probe in the mixture (top panel) to the inferred or calculated amounts of each probe in the mixture (bottom panel). The composition of each mixture is shown below the bottom panel.

First, we made an equal 3-way mixture from single positive bead sets with probes 1, 2, or 3, and analyzed it by spectral flow cytometry. Unmixing results showed relatively equal amounts of each bead type in the mixture, demonstrating that single positive “cells” could be robustly classified (Figure 6c, 6d first column). We also tested a mixture containing more of probe 1 than probes 2 and 3, and unmixing results showed relatively similar compositions compared to the known compositions (Figure 6d second column). We then investigated if various mixtures of single and double positive beads could be accurately unmixed. A mixture of single positive beads for probes 1 and 3, and double positive beads for probes 1 and 3 were analyzed, and results showed all three populations upon unmixing as expected (Figure S2). Compositions in a variety of different bead set mixtures (3-way, 2-way, double positive, and combinations of double and single positive) were measured and compared to the actual compositions of the mixtures. Overall, results demonstrate robust classification of bead (cell) type, as well as accurate estimation of the relative abundance of each bead (cell) type. (Figure 6d-compare actual to inferred heatmaps). We conclude that MuSIC probe-labeled antibodies as generated here can be used in spectral flow cytometry applications for cell type classification and proportion estimation.

## Discussion

Here we established a method to conjugate two fluorophores to an antibody in a way that enables FRET between them (if they are compatible). The use of combinations of fluorophores that exhibit FRET creates unique emission spectral signatures that can be used for multiplexing via the MuSIC approach. Antibodies are labeled with combinations of fluorophores using a modular approach combining a “linker” and DNA oligos. The linker is used to covalently attach a “docking strand” oligo to the antibody. Separate “donor” and “acceptor” strands then hybridize to the docking strand. The donor and acceptor strand oligos place the fluorophores at a specified distance from one another on the antibody. Absorbance data suggested a degree of labeling of ~9.66 linker / antibody molecules. We validated the approach using three different MuSIC probes (Cy3, Tex615, and a Cy3-Tex615 combination) attached to three separate mixtures of antibodies. MuSIC probe-labeled antibodies attached to protein A beads served as surrogate single-cells for testing via spectral flow cytometry. Spectral flow cytometry experiments demonstrated that each MuSIC probe can be uniquely differentiated by accurately determining compositions of bead mixtures.

While the focus here was using MuSIC probe labeled antibodies with spectral flow cytometry, they are also compatible in principal with spectral imaging. Several methods that increase image multiplexing capabilities use a stain/strip technique, which involves cycles of staining, imaging, and bleaching^24,26,28,43^. These methods have improved multiplexing abilities by ~10 fold over standard single-round 4-color imaging. The use of MuSIC probes is in principal compatible with the cyclic methods, which would expand the number of probes that can be used per round of imaging using spectral scanning microscopes. Current cyclic methods on average use 10 rounds of four-color imaging and our previous simulation studies suggested that ~25 MuSIC probes can be accurately unmixed^37^. Therefore, the use of MuSIC probes could allow 10 rounds of 25 color imaging, thus increasing multiplexing capabilities by roughly another six-fold. However, spectral emission scanning microscopes are certainly not as pervasive and filter-based microscopes currently. Angle-tuned emission filters for wavelength scanning may help to make such technology more accessible^44^. Such microscopes also commonly have white light lasers for tunable excitation wavelengths, and a potentially large number of channels, which would further empower multiplexing capabilities via MuSIC approaches.

To further increase fluorescent multiplexing capabilities using the MuSIC approach, additional combinations of fluorophores are needed. The FRET efficiency of a fluorophore combination is dependent on the physical distance between the two fluorophores based on the Förster radius, which is dependent on the spectral properties of the pair. Some fluorophore pairs require different distances between the two fluorophores in order to maximize FRET efficiency. This distance between the fluorophores can be varied by using different length docking strands which have varying numbers of center base pairs. Thus, we expect future solutions will use different length docking strands for different fluorophore combinations, in the march towards a larger palate of antibody-compatible MuSIC probes. Additionally, in this paper, we demonstrated unmixing of MuSIC probes using a two-laser spectral flow cytometer (488nm and 635 nm). The number of useful MuSIC probe combinations can be further increased by using a spectral flow cytometer with five excitation lasers (355 nm, 405 nm, 488nm, 561 nm, and 635nm—Cytek Aurora).

MuSIC probes have a wide variety of applications for flow cytometry, one of which is immune profiling. Immune profiling is typically performed using flow cytometry^45–47^, which has previously limited multiplexing to roughly a dozen analytes (depending on the capabilities of the instrument) as a result of spectral overlap^48,49^. Mass cytometry has been transformative for immune profiling^50–52^, but is slower than flow cytometry and is destructive so prevents further use of the cells after analysis^48^. The use of MuSIC probes for immune profiling via flow cytometry could allow for increased multiplexing for deep immune profiling on par with mass cytometry, while also being fast (more than 10,000 cells/second rather than about 1,000 as with mass cytometry^53^) and non-destructive. This could open up avenues of increased throughput for monitoring immune responses across large patient cohorts, as well as the isolation of rare cell types alone or in specified combinations that would otherwise not be possible.

We conclude that oligo-based approaches are a robust and modular way to create MuSIC probe-labeled antibodies. Future work needs to expand the MuSIC probe palette, as well as expand to larger antibody panels for flow cytometry or other spectral fluorescence applications. This would enable broader applications for advancing our understanding of microbial communities^54^ such as gut and skin microbiomes^55,56^, cancer research and clinical diagnostics, host-pathogen interactions, developmental biology, and many other areas of life science research where more highly multiplexed single and sub-cellular resolution of antibody-target readouts is informative.

## Acknowledgments

We thank Allon Klein at HMS Systems Biology for helpful discussion related to oligo-based labeling.

---

**Figure S1:**
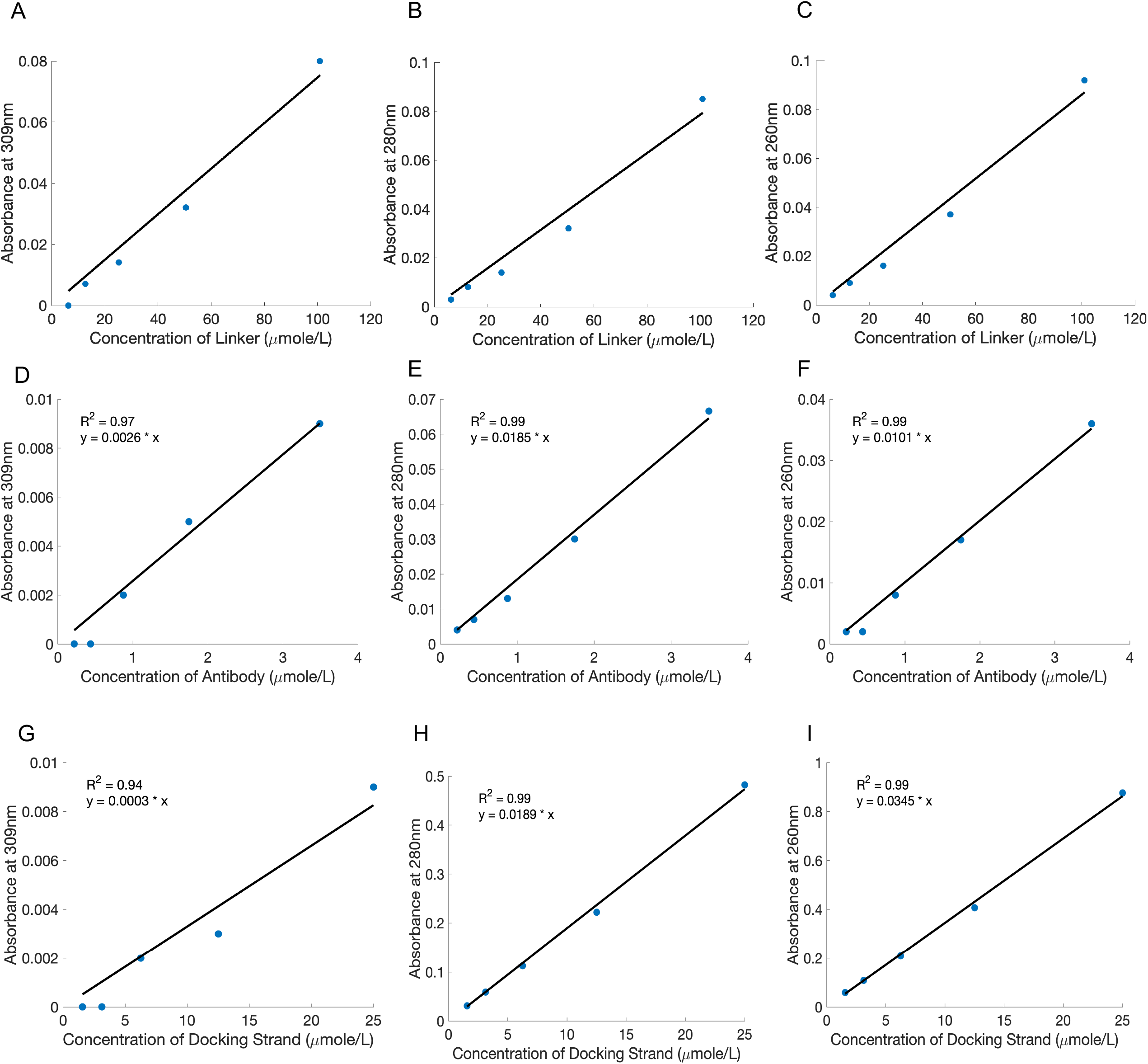
Concentration curves for the linker (A-C), antibody (D-F) and docking strand (G-I) at 309nm (A, D, G), 280nm (B, E, H), and 260nm (C, F, I). The equation of the least squares line of best fit and corresponding R^2^ values are shown on each plot. Extinction coefficients for each component at each wavelength are given from the slope of the line of best fit.

**Figure S2:**
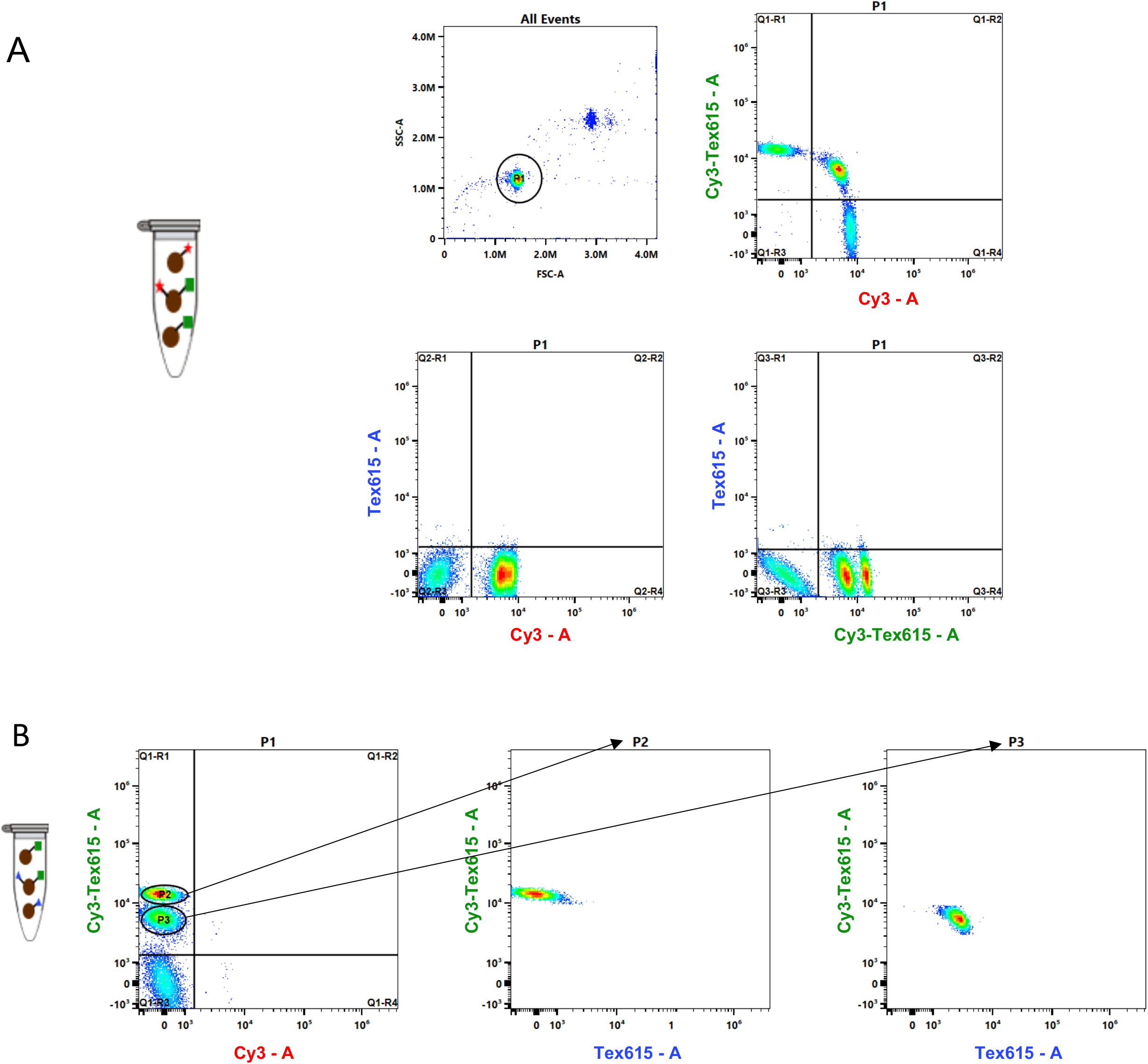
Example gating strategy on double-positive beads. (A) Gating for a mixture of beads labeled with Cy3, Cy3-Tex615, and double positive beads with both Cy3 and Cy3-Tex615. All three populations of beads are shown when plotting Cy3 vs. Cy3-Tex615 (upper right panel). (B) Gating for a mixture of beads labeled with Tex615, Cy3-Tex615, and double positive beads with both Tex615 and Cy3-Tex615. Two populations are observed in R1 when Cy3 is plotted against Cy3-tex615 (left panel). Gating on these two populations shows two distinct populations, beads labeled with just Cy3-Tex615 and double positive beads with Cy3-Tex615 and Tex615.

## References

(1) Zimmermann, T. Spectral Imaging and Linear Unmixing in Light Microscopy. In Microscopy Techniques: -/-, Rietdorf, J., Ed.; Springer Berlin Heidelberg: Berlin, Heidelberg, 2005; pp 245–265. https://doi.org/10.1007/b102216.

(2) Paolo P. Provenzano; Curtis T. Rueden; Steven M. Trier, Long Yan, Suzanne M. Ponik, David R. Inman, Patricia J. Keely, Kevin W. Eliceiri. Nonlinear Optical Imaging and Spectral-Lifetime Computational Analysis of Endogenous and Exogenous Fluorophores in Breast Cancer. J. Biomed. Opt. 2008, 13 (3), 1–11. https://doi.org/10.1117/1.2940365.

(3) Tsurui, H., Nishimura, H.; Hattori, S.; Hirose, S.; Okumura, K.; Shirai, T. Seven-Color Fluorescence Imaging of Tissue Samples Based on Fourier Spectroscopy and Singular Value Decomposition. J. Histochem. Cytochem. 2000, 48 (5), 653–662. https://doi.org/10.1177/002215540004800509.

(4) Chenying Yang, Vivian W. Hou, Leonard Y. Nelson, Eric J. Seibel. Mitigating Fluorescence Spectral Overlap in Wide-Field Endoscopic Imaging. J. Biomed. Opt. 2013, 18 (8), 1–14. https://doi.org/10.1117/1.JBO.18.8.086012.

(5) Graf, J. F.; Zavodszky, M. I. Characterizing the Heterogeneity of Tumor Tissues from Spatially Resolved Molecular Measures. PLoS ONE 2017, 12 (11), 1–20. https://doi.org/10.1371/journal.pone.0188878.

(6) Alizadeh, A. A.; Aranda, V.; Bardelli, A.; Blanpain, C.; Bock, C.; Borowski, C.; Caldas, C.; Califano, A.; Doherty, M.; Elsner, M.; Esteller, M.; Fitzgerald, R.; Korbel, J. O.; Lichter, P.; Mason, C. E.; Navin, N.; Pe’er, D.; Polyak, K.; Roberts, C. W. M.; Siu, L.; Snyder, A.; Stower, H.; Swanton, C.; Verhaak, R. G. W.; Zenklusen, J. C.; Zuber, J.; Zucman-Rossi, J. Toward Understanding and Exploiting Tumor Heterogeneity. Nat. Med. 2015, 21, 846–846.

(7) O’Connor, J. P. B.; Rose, C. J.; Waterton, J. C.; Carano, R. A. D.; Parker, G. J. M.; Jackson, A. Imaging Intratumor Heterogeneity: Role in Therapy Response, Resistance, and Clinical Outcome. Clin. Cancer Res. 2015, 21 (2), 249–257. https://doi.org/10.1158/1078-0432.CCR-14-0990.

(8) Sottoriva, A.; Spiteri, I.; Piccirillo, S. G. M.; Touloumis, A.; Collins, V. P.; Marioni, J. C.; Curtis, C.; Watts, C.; Tavaré, S. Intratumor Heterogeneity in Human Glioblastoma Reflects Cancer Evolutionary Dynamics. Proc. Natl. Acad. Sci. U. S. A. 2013, 110 (10), 4009–4014. https://doi.org/10.1073/pnas.1219747110.

(9) Gerlinger, M.; Rowan, A. J.; Horswell, S.; Math, M.; Larkin, J.; Endesfelder, D.; Gronroos, E.; Martinez, P.; Matthews, N.; Stewart, A.; Tarpey, P.; Varela, I.; Phillimore, B.; Begum, S.; McDonald, N. Q.; Butler, A.; Jones, D.; Raine, K.; Latimer, C.; Santos, C. R.; Nohadani, M.; Eklund, A. C.; Spencer-Dene, B.; Clark, G.; Pickering, L.; Stamp, G.; Gore, M.; Szallasi, Z.; Downward, J.; Futreal, P. A.; Swanton, C. Intratumor Heterogeneity and Branched Evolution Revealed by Multiregion Sequencing. N. Engl. J. Med. 2012, 366 (10), 883–892. https://doi.org/10.1056/NEJMoa1113205.

(10) Landau, D. A.; Carter, S. L.; Stojanov, P.; McKenna, A.; Stevenson, K.; Lawrence, M. S.; Sougnez, C.; Stewart, C.; Sivachenko, A.; Wang, L.; Wan, Y.; Zhang, W.; Shukla, S. A.; Vartanov, A.; Fernandes, S. M.; Saksena, G.; Cibulskis, K.; Tesar, B.; Gabriel, S.; Hacohen, N.; Meyerson, M.; Lander, E. S.; Neuberg, D.; Brown, J. R.; Getz, G.; Wu, C. J. Evolution and Impact of Subclonal Mutations in Chronic Lymphocytic Leukemia. Cell 2013, 152 (4), 714–726. https://doi.org/10.1016/j.cell.2013.01.019.

(11) Turashvili, G.; Brogi, E. Tumor Heterogeneity in Breast Cancer. Front. Med. 2017, 4, 227. https://doi.org/10.3389/fmed.2017.00227.

(12) Esposito, A.; Criscitiello, C.; Locatelli, M.; Milano, M.; Curigliano, G. Liquid Biopsies for Solid Tumors: Understanding Tumor Heterogeneity and Real Time Monitoring of Early Resistance to Targeted Therapies. Pharmacol. Ther. 2016, 157, 120–124. https://doi.org/10.1016/j.pharmthera.2015.11.007.

(13) Meldrum, C.; Doyle, M. A.; Tothill, R. W. Next-Generation Sequencing for Cancer Diagnostics: A Practical Perspective. Clin. Biochem. Rev. 2011, 32 (4), 177–195.

(14) Bagger, F. O.; Probst, V. Single Cell Sequencing in Cancer Diagnostics. In Single-cell Sequencing and Methylation: Methods and Clinical Applications, Yu, B.; Zhang, J.; Zeng, Y.; Li, L.; Wang, X.; Eds., Springer Singapore: Singapore, 2020, pp 175–193. https://doi.org/10.1007/978-981-15-4494-1_15.

(15) Müllauer, L. Next Generation Sequencing: Clinical Applications in Solid Tumours. Memo - Mag. Eur. Med. Oncol. 2017, 10 (4), 244–247. https://doi.org/10.1007/s12254-017-0361-1.

(16) Alemany, A.; Florescu, M.; Baron, C. S.; Peterson-Maduro, J.; van Oudenaarden, Whole-Organism Clone Tracing Using Single-Cell Sequencing. Nature 2018, 556 (7699), 108–112. https://doi.org/10.1038/nature25969.

(17) Wen, W.; Su, W.; Tang, H.; Le, W.; Zhang, X.; Zheng, Y.; Liu, X.; Xie, L.; Li, J.; Ye, J.; Dong, L.; Cui, X.; Miao, Y.; Wang, D.; Dong, J.; Xiao, C.; Chen, W.; Wang, H. Immune Cell Profiling of COVID-19 Patients in the Recovery Stageby Single-Cell Sequencing. Cell Discov. 2020, 6 (1), 31. https://doi.org/10.1038/s41421-020-0168-9.

(18) Gomes, T.; Teichmann, S. A.; Talavera-López, C. Immunology Driven by Large-Scale Single-Cell Sequencing. Trends Immunol. 2019, 40 (11), 1011–1021. https://doi.org/10.1016/j.it.2019.09.004.

(19) Mader, S.; Pantel, K. Liquid Biopsy: Current Status and Future Perspectives. Oncol. Res. Treat. 2017, 40 (7–8), 404–408. https://doi.org/10.1159/000478018.

(20) Chen, M.; Zhao, H. Next-Generation Sequencing in Liquid Biopsy: Cancer Screening and Early Detection. Hum. Genomics 2019, 13 (1), 34. https://doi.org/10.1186/s40246-019-0220-8.

(21) Iwahashi, N.; Sakai, K.; Noguchi, T.; Yahata, T.; Matsukawa, H.; Toujima, S.; Nishio, K.; Ino, K. Liquid Biopsy-Based Comprehensive Gene Mutation Profiling for Gynecological Cancer Using CAncer Personalized Profiling by Deep Sequencing. Sci. Rep. 2019, 9 (1), 10426. https://doi.org/10.1038/s41598-019-47030-w.

(22) Eng, C.-H. L.; Lawson, M.; Zhu, Q.; Dries, R.; Koulena, N.; Takei, Y.; Yun, J.; Cronin, C.; Karp, C.; Yuan, G.-C.; Cai, L. Transcriptome-Scale Super-Resolved Imaging in Tissues by RNA SeqFISH+. Nature 2019, 568 (7751), 235–239. https://doi.org/10.1038/s41586-019-1049-y.

(23) Rodriques, S. G.; Stickels, R. R.; Goeva, A.; Martin, C. A.; Murray, E.; Vanderburg, C. R.; Welch, J.; Chen, L. M.; Chen, F.; Macosko, E. Z. Slide-Seq: A Scalable Technology for Measuring Genome-Wide Expression at High Spatial Resolution. Science 2019, 363 (6434), 1463–1467. https://doi.org/10.1126/science.aaw1219.

(24) Gerdes, M. J.; Sevinsky, C. J.; Sood, A.; Adak, S.; Bello, M. O.; Bordwell, A.; Can, A.; Corwin, A.; Dinn, S.; Filkins, R. J.; Hollman, D.; Kamath, V.; Kaanumalle, S.; Kenny, K.; Larsen, M.; Lazare, M.; Li, Q.; Lowes, C.; McCulloch, C. C.; McDonough, E.; Montalto, M. C.; Pang, Z.; Rittscher, J.; Santamaria-Pang, A.; Sarachan, B. D.; Seel, M. L.; Seppo, A.; Shaikh, K.; Sui, Y.; Zhang, J.; Ginty, F. Highly Multiplexed Single-Cell Analysis of Formalin-Fixed, Paraffin-Embedded Cancer Tissue. Proc. Natl. Acad. Sci. 2013, 110 (29), 11982–11987. https://doi.org/10.1073/pnas.1300136110.

(25) Gut, G.; Herrmann, M. D.; Pelkmans, L. Multiplexed Protein Maps Link Subcellular Organization to Cellular States. Science 2018, 361 (6401), eaar7042–eaar7042. https://doi.org/10.1126/science.aar7042.

(26) Lin, J. R.; Fallahi-Sichani, M.; Chen, J. Y.; Sorger, P. K. Cyclic Immunofluorescence (CycIF), A Highly Multiplexed Method for Single-Cell Imaging. Curr. Protoc. Chem. Biol. 2016, 8 (4), 251–264. https://doi.org/10.1002/cpch.14.

(27) Goltsev, Y.; Samusik, N.; Kennedy-Darling, J.; Bhate, S.; Hale, M.; Vazquez, G.; Black, S.; Nolan, G. P. Deep Profiling of Mouse Splenic Architecture with CODEX Multiplexed Imaging. Cell 2018, 174 (4), 968-981.e15. https://doi.org/10.1016/j.cell.2018.07.010.

(28) Lin, J. R.; Izar, B.; Wang, S.; Yapp, C.; Mei, S.; Shah, P. M.; Santagata, S.; Sorger, P. K. Highly Multiplexed Immunofluorescence Imaging of Human Tissues and Tumors Using T-CyCIF and Conventional Optical Microscopes. eLife 2018, 7, 1–46. https://doi.org/10.7554/eLife.31657.

(29) Giesen, C.; Wang, H. A. O.; Schapiro, D.; Zivanovic, N.; Jacobs, A.; Hattendorf, B.; Schüffler, P. J.; Grolimund, D.; Buhmann, J. M.; Brandt, S.; Varga, Z.; Wild, P. J.; Günther, D.; Bodenmiller, B. Highly Multiplexed Imaging of Tumor Tissues with Subcellular Resolution by Mass Cytometry. Nat. Methods 2014, 11 (4), 417–422. https://doi.org/10.1038/nmeth.2869.

(30) Angelo, M.; Bendall, S. C.; Finck, R.; Hale, M. B.; Hitzman, C.; Borowsky, A. D.; Levenson, R. M.; Lowe, J. B.; Liu, S. D.; Natkunam, Y.; Nolan, G. P. Multiplexed Ion Beam Imaing (MIBI) of Human Breast Tumors. Nat. Med. 2014, 20 (4), 436–442. https://doi.org/10.1038/nm.3488.Multiplexed.

(31) Haraguchi, T.; Shimi, T.; Koujin, T.; Hashiguchi, N.; Hiraoka, Y. Spectral Imaging Fluorescence Microscopy. Genes Cells 2002, 7 (9), 881–887. https://doi.org/10.1046/j.1365-2443.2002.00575.x.

(32) Lu, G.; Fei, B. Medical Hyperspectral Imaging: A Review. J. Biomed. Opt. 2014, 19 (1), 010901–010901. https://doi.org/10.1117/1.jbo.19.1.010901.

(33) Martin, M. E.; Wabuyele, M. B.; Chen, K.; Kasili, P.; Panjehpour, M.; Phan, M.; Overholt, B.; Cunningham, G.; Wilson, D.; DeNovo, R. C.; Vo-Dinh, T. Development of an Advanced Hyperspectral Imaging (HSI) System with Applications for Cancer Detection. Ann. Biomed. Eng. 2006, 34 (6), 1061–1068. https://doi.org/10.1007/s10439-006-9121-9.

(34) Akbari, H.; Halig, L.; Schuster, D. M.; Fei, B.; Osunkoya, A.; Master, V.; Nieh, P.; Chen, G. Hyperspectral Imaging and Quantitative Analysis for Prostate Cancer Detection. J. Biomed. Opt. 2012, 17 (7), 1–11.

(35) Valm, A. M.; Mark Welch, J. L., Borisy, G. G. CLASI-FISH: Principles of Combinatorial Labeling and Spectral Imaging. Spec. Issue Fluoresc. Situ Hybrid. FISH 2012, 35 (8), 496–502. https://doi.org/10.1016/j.syapm.2012.03.004.

(36) Niehörster, T.; Löschberger, A.; Gregor, I.; Krämer, B.; Rahn, H.-J.; Patting, M.; Koberling, F.; Enderlein, J.; Sauer, M. Multi-Target Spectrally Resolved Fluorescence Lifetime Imaging Microscopy. Nat. Methods 2016, 13, 257–257.

(37) Holzapfel, H. Y.; Stern, A. D.; Bouhaddou, M.; Anglin, C. M.; Putur, D.; Comer, S.; Birtwistle, M. R. Fluorescence Multiplexing with Spectral Imaging and Combinatorics. ACS Comb. Sci. 2018, 20 (11), 653–659. https://doi.org/10.1021/acscombsci.8b00101.

(38) Clapp, A. R.; Medintz, I. L.; Mattoussi, H. Förster Resonance Energy Transfer Investigations Using Quantum-Dot Fluorophores. ChemPhysChem 2006, 7 (1), 47–57. https://doi.org/10.1002/cphc.200500217.

(39) Holzapfel, H. Y.; Birtwistle, M. R. Creating Complex Fluorophore Spectra on Antibodies Through Combinatorial Labeling. Transl. Sci. 2016, 2 (3), e03.

(40) Gong, H.; Holcomb, I.; Ooi, A.; Wang, X.; Majonis, D.; Unger, M. A.; Ramakrishnan, R. Simple Method To Prepare Oligonucleotide-Conjugated Antibodies and Its Application in Multiplex Protein Detection in Single Cells. Bioconjug. Chem. 2016, 27 (1), 217–225. https://doi.org/10.1021/acs.bioconjchem.5b00613.

(41) Varn, F. S.; Tafe, L. J.; Amos, C. I.; Cheng, C. Computational Immune Profiling in Lung Adenocarcinoma Reveals Reproducible Prognostic Associations with Implications for Immunotherapy. OncoImmunology 2018, 7 (6), e1431084. https://doi.org/10.1080/2162402X.2018.1431084.

(42) Goswami, S.; Walle, T.; Cornish, A. E.; Basu, S.; Anandhan, S.; Fernandez, I.; Vence, L.; Blando, J.; Zhao, H.; Yadav, S. S.; Ott, M.; Kong, L. Y.; Heimberger, A. B.; de Groot, J.; Sepesi, B.; Overman, M.; Kopetz, S.; Allison, J. P.; Pe’er, D.; Sharma, P. Immune Profiling of Human Tumors Identifies CD73 as a Combinatorial Target in Glioblastoma. Nat. Med. 2020, 26 (1), 39–46. https://doi.org/10.1038/s41591-019-0694-x.

(43) Trindade, C. J.; McDonough, E.; Hanson, J.; Walter Rodriguez, B.; Roper, N.; Gasmi, B.; Roque, C.; Gebregziabher, M.; Ylaya, K.; Fetsch, P.; Abdul Sater, H.; Ginty, F.; Hewitt, S. M.; Thomas, A. Utilization of Novel Highly Multiplexed Immunofluorescence Microscopy Technology to Understand Immunological Tumor Microenvironments in Small Cell Lung Carcinoma Patients Receiving Combination PD-L1 and PARP Inhibition Therapy. J. Clin. Oncol. 2019, 37 (15_suppl), e14289–e14289. https://doi.org/10.1200/JCO.2019.37.15_suppl.e14289.

(44) Yu, K.; Liu, Y.; Yin, J.; Bao, J. A Novel Angle-Tuned Thin Film Filter with Low Angle Sensitivity. Opt. Laser Technol. 2015, 68, 141–145. https://doi.org/10.1016/j.optlastec.2014.11.022.

(45) Daud, A. I.; Loo, K.; Pauli, M. L.; Sanchez-Rodriguez, R.; Sandoval, P. M.; Taravati, K.; Tsai, K.; Nosrati, A.; Nardo, L.; Alvarado, M. D.; Algazi, A. P.; Pampaloni, M. H.; Lobach, I. V.; Hwang, J.; Pierce, R. H.; Gratz, I. K.; Krummel, M. F.; Rosenblum, M. D. Tumor Immune Profiling Predicts Response to Anti–PD-1 Therapy in Human Melanoma. J. Clin. Invest. 2016, 126 (9), 3447–3452. https://doi.org/10.1172/JCI87324.

(46) Koutsakos, M.; Sekiya, T.; Chua, B. Y.; Nguyen, T. H. O.; Wheatley, A. K.; Juno, J. A.; Ohno, M.; Nomura, N.; Ohara, Y.; Nishimura, T.; Endo, M.; Suzuki, S.; Ishigaki, H.; Nakayama, M.; Nguyen, C. T.; Itoh, Y.; Shingai, M.; Ogasawara, K.; Kino, Y.; Kent, S. J.; Jackson, D. C.; Brown, L. E.; Kida, H.; Kedzierska, K. Immune Profiling of Influenza-Specific B-and T-Cell Responses in Macaques Using Flow Cytometry-Based Assays. Immunol. Cell Biol. 2020, n/a (n/a). https://doi.org/10.1111/imcb.12383.

(47) Vacchi, E.; Burrello, J.; Di Silvestre, D.; Burrello, A.; Bolis, S.; Mauri, P.; Vassalli, G.; Cereda, C. W.; Farina, C.; Barile, L.; Kaelin-Lang, A.; Melli, G. Immune Profiling of Plasma-Derived Extracellular Vesicles Identifies Parkinson Disease. Neurol. - Neuroimmunol. Neuroinflammation 2020, 7 (6), e866. https://doi.org/10.1212/NXI.0000000000000866.

(48) Landhuis, E. S Ingle-Cell Approaches to Immune Profiling. Nature 2018, 557 (7706), 595—597. https://doi.org/10.1038/d41586-018-05214-w.

(49) Spitzer, M. H.; Nolan, G. P. Mass Cytometry: Single Cells, Many Features. Cell 2016, 165 (4), 780–791. https://doi.org/10.1016/j.cell.2016.04.019.

(50) Bengsch, B.; Ohtani, T.; Herati, R. S.; Bovenschen, N.; Chang, K.-M.; Wherry, E. J. Deep Immune Profiling by Mass Cytometry Links Human T and NK Cell Differentiation and Cytotoxic Molecule Expression Patterns. Mass Cytom. Methods Appl. 2018, 453, 3–10. https://doi.org/10.1016/j.jim.2017.03.009.

(51) Böttcher, C.; Fernández-Zapata, C.; Schlickeiser, S.; Kunkel, D.; Schulz, A. R.; Mei, H. E.; Weidinger, C.; Gieß, R. M., Asseyer, S.; Siegmund, B.; Paul, F.; Ruprecht, K.; Priller, J. Multi-Parameter Immune Profiling of Peripheral Blood Mononuclear Cells by Multiplexed Single-Cell Mass Cytometry in Patients with Early Multiple Sclerosis. Sci. Rep. 2019, 9 (1), 19471. https://doi.org/10.1038/s41598-019-55852-x.

(52) Wang, W.; Su, B.; Pang, L.; Qiao, L.; Feng, Y.; Ouyang, Y.; Guo, X.; Shi, H.; Wei, F.; Su, X.; Yin, J.; Jin, R.; Chen, D. High-Dimensional Immune Profiling by Mass Cytometry Revealed Immunosuppression and Dysfunction of Immunity in COVID-19 Patients. Cell. Mol. Immunol. 2020, 17 (6), 650–652. https://doi.org/10.1038/s41423-020-0447-2.

(53) Li, L.; Yan, S.; Lin, B.; Shi, Q.; Lu, Y. Chapter Eight - Single-Cell Proteomics for Cancer Immunotherapy. In Advances in Cancer Research, Broome, A.-M.; Ed., Academic Press, 2018, Vol. 139, pp 185–207. https://doi.org/10.1016/bs.acr.2018.04.006.

(54) Amann, R.; Fuchs, B. M. Single-Cell Identification in Microbial Communities by Improved Fluorescence in Situ Hybridization Techniques. Nat. Rev. Microbiol. 2008, 6 (5), 339–348. https://doi.org/10.1038/nrmicro1888.

(55) Tropini, C.; Earle, K. A.; Huang, K. C.; Sonnenburg, J. L. The Gut Microbiome: Connecting Spatial Organization to Function. Cell Host Microbe 2017, 21 (4), 433–442. https://doi.org/10.1016/j.chom.2017.03.010.

(56) Brandwein, M.; Steinberg, D.; Meshner, S. Microbial Biofilms and the Human Skin Microbiome. Npj Biofilms Microbiomes 2016, 2 (1), 3. https://doi.org/10.1038/s41522-016-0004-z.

